# Ovariectomy as a Management Technique in Suburban Deer Populations

**DOI:** 10.1101/2020.07.22.216481

**Authors:** Anthony J. DeNicola, Vickie L. DeNicola

**Affiliations:** White Buffalo Incorporated, 26 Davison Road, Moodus, CT 06469, USA

**Keywords:** deer management, human-wildlife conflicts, nonlethal, *Odocoileus* spp, ovariectomy, population management

## Abstract

Overabundant suburban deer (*Odocoileus* spp.) are a source of human-wildlife conflict in many communities throughout the United States. Deer-vehicle collisions, tick-borne pathogens, impacts on local vegetation, and other negative interactions are the typical reasons cited for initiating a deer management program. Social attitudes, legal constraints, and perceived safety concerns lead many communities to examine nonlethal management options. Surgical sterilization is currently the only nonlethal method available to permanently sterilize females with a single treatment. There are limited data demonstrating methods and outcomes in management programs that sterilize a high percentage (>90%) of the local population, particularly regarding the impact of immigration on non-isolated populations. We present data from 6 surgical sterilization sites with geographically open populations in California, Maryland, Michigan, New York, Ohio, and Virginia, USA. From 2012–2020, we sterilized 493 deer primarily via ovariectomy. We conducted annual or periodic population estimates using camera surveys, road-based distance sampling, and intensive field observations to assess population trends. Initial densities ranged from about 6–63 deer/km^2^. Study sites ranged from 1.2 km^2^ to 16.5 km^2^, and initial populations ranged from ~47 to 169 individuals. For our 6 study sites, we noted an average reduction in deer abundance of approximately 25% (range: 16.2%–36.2%) from Year 1 to Year 2. Four years after the first treatment, at monitored sites (*n* = 4) using this management method, we noted an average total population reduction of about 45% (range: 28%–56%). During the first year, the average cost per deer handled was $1,221 (range: $864–$1,998). These projects demonstrate that significant reductions in local deer densities using high percentage surgical sterilization programs can be achieved in non-insular locations. Sustained sterilization efforts are necessary, as is the case with all deer management programs in open landscapes.

---

In many urban and suburban situations, lethal management techniques for deer (*Odocoileus* spp.) such as controlled hunting and sharpshooting are not feasible or practical because of legal, social, public safety, or economic considerations (Williams et al. 2013, Kilpatrick et al. 2014). As a result, various non-lethal management techniques such as trap and relocation or fertility control have been explored (DeNicola et al. 2000). Fertility control options have been prioritized because trap and relocation has been demonstrated to be stressful and result in unacceptably high post-release mortality (Beringer et al. 2002).

Extensive research has been conducted on injectable or implantable wildlife fertility control agents for nearly 5 decades, with numerous studies conducted in the 1970s and 1980s (Harder and Peterle 1974, Bell and Peterle 1975, Matschke 1977, Kirkpatrick and Turner 1985, Plotka and Seal 1989). Recent research has transitioned focus to vaccines and surgical sterilization as the most promising approaches (Patten et al. 2007). Given the scope of research, there has been relatively little improvement in the function of vaccines tested in the late-1980s on horses (Liu et al. 1989, Kirkpatrick et al. 1990) or deer in the 1990s (Turner et al. 1992, McShea et al. 1997, Miller et al. 1997, Miller et al. 2000). For long-lived species, at least one booster is necessary to predictably maintain >90% effectiveness for >2 years, even for vaccines formulated as a single dose (Gionfriddo et al. 2011, Rutberg et al. 2013a, Roelle et al. 2017).

Wildlife professionals have discussed the potential of fertility control as a wildlife management technique for as long as efficacy trials have been conducted (McCullough 1987, Bomford 1992, Garrott 1995, Warren 1995, Miller et al. 1998). Most studies have focused on delivery methods and treatment efficacy, with very few assessments of management outcomes. In addition, research to date has primarily described the limitations of fertility control versus its potential to effectively manage deer overabundance in geographically open environments. In fact, several studies have concluded that for fertility control to be successful, its use must be limited to small, insular, or fenced areas (Seagle and Close 1996, Merrill et al. 2006, Boulanger et al. 2012, Boulanger and Curtis 2016). The conclusion that the method is not viable in open populations is founded on population-level field studies conducted at the Cornell University Campus, Ithaca, New York, USA (Boulanger and Curtis 2016), Fripp Island, South Carolina, USA (Rutberg et al. 2013b), National Institute of Standards and Technology, Gaithersburg, Maryland, USA (Rutberg and Naugle 2008), and Fire Island National Seashore, Patchogue, New York, USA (Naugle et al. 2002), as well as various modeling efforts (Seagle and Close 1996, Merrill et al. 2006). One modeling effort projected a reduced local population using surgical sterilization at Cumberland Island, Georgia, USA (Boone and Wiegert 1994).

Conclusions on the potential management benefits of fertility control have been based on compromised methods and incorrect assumptions regarding immigration rates. In the Cornell University study that used tubal ligation as the surgical method, it was concluded that surgical sterilization would not work in geographically open environments without a lethal component (Boulanger and Curtis 2016). In this study, adult female and fawn numbers declined significantly, reflecting an effective program, but male abundance increased 9-fold, preventing an overall population decline. Given the sterilization method used caused extended estrous cycles, males were potentially attracted to females during later estrus periods. Antlered males may also have sought refuge in the non-hunted study site, given the earn-a-buck hunting program that occurred around the site (Williams et al. 2008, Boulanger et al. 2014). Another study found limited immigration and no increase of adult males due to sterilization of adult females via tubal ligation in an open suburban environment without proximate hunting (MacLean et al. 2006).

The first assessment of surgical sterilization on deer was conducted in the early 1990s (Frank and Sajdak 1993) and additional research on surgical methods occurred in the 2000s (MacLean et al. 2006, Boulanger et al. 2012). These efforts were initiated in recognition that immunocontraceptive vaccines were limited in their long-term effectiveness, and an alternative was needed in areas where treating the same animal more than once was too difficult or costly. As a result, we decided to conduct additional fertility control research, using surgical sterilization via ovariectomy, to reduce population abundance while eliminating breeding behavior associated with tubal ligations. Our interest was to identify a method that could be used as an infertility treatment and have population-level impacts on white-tailed deer (*O. virginianus*). Our research focused on the potential of surgical sterilization by ovariectomy of female deer as a management tool in geographically open suburban environments and suburban environments with security fences permeable to deer. Given the strong philopatry of female white-tailed deer, immigration could be minimal (Porter et al. 2004). We selected research locations based on fundamental needs for extensive access and approachable deer (McCullough 1987, Garrott 1995, Rudolph et al. 2000). Most researchers have concluded that 60–80% or greater of a deer population needs to be rendered infertile to stabilize or begin to reduce a local population (McCullough 1987, Hone 1992, Garrott 1995, Grund 2011, Boulanger et al. 2012). Our objectives in each designated study area were to 1) capture and permanently sterilize >95% of females; 2) assess immigration rates, mortality, and changes in population abundance; and 3) assess effort.

## STUDY AREAS

We applied surgical sterilization treatments to free-ranging deer on 6 study sites. White-tailed deer were the target species at every site except The Villages Golf and Country Club (VGCC), where we captured and treated black-tailed deer (*O. hemionus columbianus*). Deer capture efforts were conducted between August and March from 2012–2020 to minimize stress on deer and maximize baiting success. The justification for attempting to reduce deer abundance included increasing deer-vehicle collisions, landscape damage, forest diversity impacts, or tick-borne infection concerns. Decision-makers at each location opted to use a non-lethal approach because of either social or legal constraints.

Our study sites were typified by a mixture of developed urban or suburban landscapes with intermingled wetlands or other small to moderate open spaces. Regional differences in climate among our various study sites affected the local natural plant communities. Temperate, mesic hardwood forests predominantly composed the open areas at 5 study sites. Xeric shrublands with dry oak savanna surrounded VGCC. Except for the National Institutes of Health (NIH) campus, the sites were primarily single-family residential communities with property sizes ranging from 0.4–2.0 ha with some larger private properties interspersed, such as golf courses. Only the NIH campus and the VGCC community had security fencing. Neither fence prevented deer from entering or exiting with documented emigration from both locations and annual immigration at NIH. The other study sites were not isolated from the surrounding areas that contained resident deer populations. The study sites did not permit hunting because of safety concerns associated with high-density development and significant human activity. Hunting or deer management activities occurred in communities adjacent to 4 of our study sites.

The study site in Ann Arbor, Michigan, USA (N42.2709, W83.7263; AA) consisted of 4.3 km^2^, and most of its land area was covered by single-family residential communities surrounded by wooded corridors (Fig. 1). The community opted to incorporate culling into the broader program scope because of the disparity in development patterns. Sharpshooting occurred adjacent to the northern and eastern borders of the study site. Sterilized deer were not culled during sharpshooting operations. The City’s management goal was to reduce deer densities to the point where 75% of surveyed residents reported satisfaction with the deer abundance.

The Clifton neighborhood of Cincinnati, Ohio, USA (N39.1531, W84.5248; CLIF), was located in the north-central section of the city and the 2.4 km^2^ study site contained Mt. Storm, Rawson Woods, and Edgewood Grove parks, as well as the surrounding neighborhoods consisting of single-family residential communities and apartment complexes. The parks were generally undeveloped and included steep wooded hillsides and some mowed open space. Because deer were not limited to the parks under consideration, we delineated the study area as depicted in Fig. 2. Deer densities had reached a level that was considered incompatible with the Cincinnati Park Board’s goal of a healthy park ecosystem.

Fairfax City, Virginia, USA (N38.8462, W77.3064; FC) was located in the western suburbs of Washington, D.C., and is ~16.5 km^2^ (Fig. 3). The area consisted of single-family residential communities, apartment complexes, and commercial-use properties with wooded corridors. The use of bait for capture or camera survey population estimates was not permitted during the first year. After we demonstrated that baiting would be instrumental to the project’s success and improved accuracy of camera surveys, Virginia Department of Game and Inland Fisheries (VA DGIF) amended the research permit to allow the use of bait.

The NIH site (N38.9992, W77.1025) was an office park complex located in Bethesda, Maryland, USA (Fig. 4). The study area was 1.2 km^2^ and consisted of more than 95 office buildings in a campus-like environment. The site contained small natural wooded and landscaped areas interspersed with the dense commercial development. Landscape damage and deer-vehicle collisions were of concern.

The Village of Cayuga Heights, New York, USA (N42.4687, W76.4856; VCH) was located adjacent to and contiguous with the City of Ithaca and contained 4.6 km^2^ (Fig. 5). The area consisted of single-family residential communities, apartment complexes, and commercial-use properties with wooded corridors. At the start of this study, New York State law prevented discharge of a weapon such as a bow, muzzleloader, or firearm within 152 m of an occupied dwelling without owner permission. As a result, and given local opposition to a lethal management program, there were no legal locations to cull deer in VCH. Therefore, elected officials voted to implement a surgical sterilization initiative starting in 2012, given the abundance of deer and associated conflicts. In 2014, the law changed reducing archery discharge to 76 m, which permitted a cull that was conducted from 2015–2017. The community leaders decided to incorporate archery culling to accelerate the population decline. The State of New York then prohibited baiting within 91 m of a public road in 2017, and only 2 properties remained legal given the high road density. As a result, VCH decided to transition to a remote immobilization program followed by euthanasia from 2018–2019 to obtain even lower deer densities. We included data and discussion on these subsequent culling activities to allow for immigration insights as the population declined. However, we only included data collected prior to culling in our analysis.

The VGCC was located in San Jose, California, USA (N37.2878, W121.7466) and consisted of a 2.7 km^2^ private community with 2,536 homes and an 18-hole golf course (Fig. 6). Landscape damage and replacement costs exceeded $250,000 annually because of deer impacts (B. Barncord, personal communication), and the community elected to execute a non-lethal management program as a result.

## METHODS

Licensed and trained veterinarians conducted all surgical procedures. White Buffalo Inc. employees, trained by a wildlife veterinarian in humane capture and chemical restraint methods, conducted all capture efforts. We performed all field procedures under Michigan Department of Natural Resources Scientific Collectors Permit #1600, Ohio Department of Natural Resources Scientific Collection Permit #18-48, VA DGIF Scientific Collection Permit #058581, Maryland Department of Natural Resources Deer Cooperator Permit 55832-NIH-2014-2019, and NIH Office of Research Services Animal Study Proposal Animal Care and Use Committee #ORS-54, New York State Department of Environmental Conservation License to Collect and Possess #2558, and California Department of Fish and Wildlife Scientific Collection Permit #SC-012522.

### Deer Capture

We conducted deer capture and sterilization activities between August and March from 2012– 2020. We immobilized female white-tailed deer of all age classes using projectors with 2-ml transmitter darts (Pneu-Dart, Inc., Williamsport, PA, USA) to administer tiletamine/zolazepam (4.4 mg/kg) and xylazine hydrochloride (2.2 mg/kg; Kilpatrick et al. 1997). We approached deer in a vehicle on public roadways and on private roadways and properties where permission was granted. In AA, FC, and VCH, capture professionals were accompanied by a police officer during mobile darting efforts (Evans et al. 2016). We also darted deer over 20–25 kg of whole-kernel corn bait in the late afternoon and early evening where state regulations permitted. Ten minutes after a dart was deployed, we located deer via radio telemetry, if not visibly recumbent. In VCH, we also captured deer with drop nets during the first year. We chemically immobilized adult deer under drop nets with an intramuscular injection of 400 mg ketamine hydrochloride (HCl) and 150 mg xylazine HCl <2 minutes after capture. We administered 300 mg ketamine HCl and 100 mg xylazine HCl to fawns. We fitted all deer captured with ear tags for individual identification (Destron Fearing Duflex X-Large Cattle Tag, Dallas, TX, USA) and the backplate of each labeled “Call Before Consumption” and a phone number. All individuals were transported to temporary surgical facilities, except for NIH and 2 years in CLIF, where we used a veterinary facility. We followed previous deer handling, tagging, very high frequency (VHF) radio-collaring, and transport methods (Evans et al. 2016).

### Surgical Sterilization Procedure

Initially, we conducted bilateral ovariectomies using a combination of clamping, electrocautery, and excision for removal and coagulation to prevent hemorrhage (Evans et al. 2016). We transitioned to a vessel sealer in fall 2016 to expedite the procedure and minimize risks associated with post-surgical bleeding (Valleylab LigaSure, Medtronic, Minneapolis, MN, USA). We conducted fallopian tubal ligectomies (*n* = 26) only when deemed necessary by veterinarians due to extensive adhesions that complicated the extraction of the ovaries or during late gestation.

We also followed previous post-surgical release and monitoring procedures (Evans et al. 2016). The one exception was that we relocated 30 deer outside the fence at VGCC to adjacent VGCC-owned property to assess whether these deer would return or remain outside and increase the rate of population decline. To differentiate these relocated deer from the rest of the herd, they received one yellow and one white extra-large ear tag.

### Effort

We recorded start and stop times to assess the person-hours required for capture, handling, transport, and sterilization surgery. Trained biologists and volunteers conducted capture, handling and transportation to and from surgical locations. For each study site, we divided the number of person-hours devoted to capture, handling, and transport by the number of deer processed to derive an estimate of effort per deer. Each surgical team consisted of 2 people, including trained and licensed veterinarians and veterinary technicians. We had two veterinary teams performing surgeries during the first year at all project sites. Each capture night’s surgical effort was recorded starting when the first deer was darted and ending when the last deer was recovered. We reported person-hours per deer for surgical effort.

### Population Estimates

We conducted population monitoring using camera data from baited locations, driving distance transects, and intensive field observations. We noted immigrants, antlered males, fawns, and both tagged and untagged females. We defined immigrants as untagged deer that were not identified the previous year.

At AA, CLIF and FC, we conducted baited camera surveys using tagged deer to estimate deer abundance (Jacobson et al. 1997, Koerth et al. 1997, Curtis et al. 2009). We used game cameras (D-80 White Flash Trail Cameras, Moultrie, Alabaster, AL, USA) set on motion activated single shot with a 5-minute delay to optimize capture rates relative to photo storage. We used camera coverage of ~1/0.61 km^2^ with one camera placed in each block. Project staff pre-baited sites for at least 7 days. We placed each camera 5–6 m from the center of bait, elevated 0.6 m, oriented north, and left for about 2 weeks. We retrieved cameras once we obtained about 300 photos from each baited location. There were some situations where we retrieved cameras with <300 photos due to low deer activity. In the first year of FC, baiting was not permitted. To conduct this unbaited camera survey, we divided FC into 12 quadrants and allocated 1 camera to each quadrant. We placed cameras on public property in each quadrant at locations proximate to heavily traveled deer trails and pinch points through which deer were forced to pass.

Each image was studied, all legible ear tag numbers were documented, and images of insufficient quality were excluded due to lens flares or poor image quality. We recorded the total number of deer, tagged and untagged individuals, unidentifiable tagged deer, identifiable and unidentifiable antlered males, and untagged fawns in each photo. If a tagged deer was not observed on camera or directly for 2 years, we eliminated the individual from that year’s population estimate.

We entered the number of tagged, untagged, and tagged but unidentifiable observations from each photograph into Program NOREMARK (White 1996). Our tagged individuals were adult females, with occasional female fawns and incidental male fawns. We used the Bowden estimator that assumes sightability is not influenced by tag status, many animals are sighted repeatedly, and the population is closed (Bowden 1993, Bowden and Kufel 1995). Our study areas did not have fully closed populations, but there was likely minimal outside exchange given deer philopatry, relatively small suburban home ranges, and the camera survey’s short duration. The resulting population estimate and confidence intervals for this method were for antlerless deer (95% CI). We used the buck:doe ratio (BDR) method to estimate the number of individual antlered males (Jacobson et al. 1997). We derived a total estimate using the BDR method when we did not utilize other methods. We then added the number of individual antlered males identified using the BDR method (BDR M) to the antlerless estimates from Program NOREMARK to obtain a population estimate noted as NR-BDR. For a second estimate, we used camera data to determine the ratio of tagged to untagged adult females photographed to estimate the total number of adult females and fawns, assuming equal sightability (Eberhardt 1969). We then added BDR M to the photographic ratio estimates of adult females and fawns to obtain a population estimate, noted as LPE-BDR method. Finally, we used a census approach to derive the population size. We determined the number of adult females and fawns, given the high percentage capture status, intensive field observations (e.g., capture observations, bait site camera data, population estimate data), known mortality and known dispersal data. This census also included BDR M and field observations. We then averaged the methods used to derive a final estimate. We also used field observations and photographic data to determine the percentage of females sterilized after each annual capture session, antlerless immigration, and the fawn:adult female ratio.

In VCH, the municipality conducted independent baited camera surveys and population estimates using NOREMARK from 2013–2016, after capture efforts were completed in 2013– 2014), and prior to culling efforts in 2015–2017 (Curtis et al. 2013, 2014; Curtis and Ashdown 2015, 2016, 2017). Two NOREMARK estimates were made, one using all deer identifiable in photographs and a second using the maximum number of marked deer potentially available in the study area. These two estimates were averaged to obtain a final estimate. In 2018, we used a census through field observations and camera data to estimate the population.

In VGCC, we established a non-overlapping 14.6-km driving transect and surveyed 3–5 times, either during the morning or evening, from 7–8 September 2011 and 9–11 October 2012. We conducted surveys starting 2 hours before sunset and 1 hour after sunrise. While driving 15– 20 kph in a golf cart, spotters searched their respective side of the road. Upon sighting deer, we recorded the number in each social group, age, and sex of the individuals, whether deer were tagged or untagged, and we measured the perpendicular distance the group’s center using a rangefinder. These data were entered into the software program Distance (Version 6.0; Thomas et al. 2010) to determine a population estimate and a 95% confidence interval.

In VGCC, beginning in fall 2014, 10–12 volunteers were assigned to separate neighborhoods and recorded observations of every deer they encountered, including tag numbers and antler patterns. We tallied the number and age class of individual animals observed as an estimate of the population because the deer were very visible. Observers were active for 2–3 hours before sunset for 5 days.

Given the small area and relatively open landscape at NIH, we walked the available habitat areas 2–3 hours before sunset and recorded tagged individuals, males based on antler configuration, and untagged antlerless animals, noting any fawns. We walked the facility twice during each estimate and drove intensively over the 2-day field operation. We used individual animals observed as a census of the population because the deer were highly visible.

## RESULTS

### Capture and Surgical Sterilization

We handled 570 deer, including 493 surgical sterilization procedures, incidental male captures (*n* = 61), recaptures (*n* = 10), and animals euthanized due to pre-existing conditions (*n* = 2) or capture related injuries (*n* = 4) at the 6 study sites (Table 1). We experienced 15 capture and handling related mortalities, 4 were individuals euthanized prior to surgery due to capture related injuries, and 11 were post-surgical mortalities, resulting in a 2.6% mortality rate. During one field session at VCH, we captured 29 incidental males under the drop net. Otherwise, incidental males accounted for 32 captures across all other sites and sessions. We observed 6 failed surgeries where treated females were later detected with fawns. When surgeries failed, single fawns were recruited, and in all cases, animals were recaptured and retreated. One deer with a failed surgery in VCH had an ovary remaining that was extracted upon recapture. Veterinarians were unable to locate the uterus in one deer in AA, a suspected previous prolapse, and the individual was later recaptured and successfully sterilized. We documented four surgical failures in CLIF due to a deficient surgical method that left remnant ovarian tissue. The entire ovarian capsule was not removed, allowing for regrowth of tissue. At CLIF, we recaptured 2 females to confirm lactational status and check for failed surgeries, and at VCH, 2 treated females were incidentally recaptured with the drop net. We treated an average of 84% (range: 31–100%) of the females in the first year, and 94% (range: 89–100%) in the second year at each project site.

**Table 1.** White-tailed deer surgical sterilization study site locations, field implementation dates, quantity of females sterilized, capture and handling related mortalities, and the associated number of deer captured during each field session from 2012–2020. AA = Ann Arbor, Michigan, USA; CLIF = Clifton area, Cincinnati, Ohio, USA; FC = City of Fairfax, Virginia, USA; NIH = National Institutes of Health, Bethesda, Maryland, USA; VCH = Village of Cayuga Heights, New York, USA; VGCC = Villages Golf and Country Club, San Jose, California, USA

| Study sites and<br>program dates | Females<br>sterilized | Mortality | Captured<sup>a</sup> |
| --- | --- | --- | --- |
| AA |  |  |  |
| 22–29 Jan 2017 | 49 | 0 | 52 |
| 2–6 Jan 2018 | 10 | 0 | 14 |
| 28–30 Nov 2018 | 1 | 0 | 1 |
| 22 Jan–20 Feb 2020 | 0 | 0 | 0 |
| CLIF |  |  |  |
| 1–7 Dec 2015 | 41 | 2 | 44 |
| 16–19 Jan 2017 | 10 | 0 | 10 |
| 8–14 Jan 2018 | 8 | 0 | 11 |
| 24–27 Aug 2018 | 4 | 0 | 6 |
| 24–28 Jan 2019 | 7 | 0 | 9 |
| 1 Nov 2019–8 Feb 2020 | 9 | 1 | 10 |
| FC |  |  |  |
| 31 Jan–6 Feb 2014 | 18 | 0 | 21 |
| 26–31 Jan 2015 | 18 | 0 | 20 |
| 14–16 Dec 2015 | 6 | 1 | 8 |
| 6–7 Dec 2016 | 5 | 0 | 5 |
| 16–18 Jan 2018 | 4 | 0 | 4 |
| NIH |  |  |  |
| 14–16 Dec 2014 | 24 | 2 | 27 |
| 2 Dec 2015 | 5 | 1 | 5 |
| 3 Dec 2016 | 8 | 1 | 8 |
| 4 Dec 2017 | 2 | 0 | 2 |
| 8 Dec 2018 | 3 | 0 | 3 |
| VCH |  |  |  |
| 1–15 Dec 2012 | 137 | 3 | 172 |
| 2–6 Dec 2013 | 11 | 0 | 13 |
| 26–27 Mar 2016 | 5 | 0 | 8 |
| VGCC |  |  |  |
| 25 Jan–4 Feb 2013 | 99 | 4 | 107 |
| 8–12 Oct 2013 | 9 | 0 | 10 |
| Total | 493 | 15 | 570 |

<sup>a</sup>Includes females sterilized, recaptures, incidental male captures, and capture/pre-existing conditions related euthanasia.

### Surgical and Capture Effort and Cost

Effort by field staff in Year 1 averaged 4.9 hours/deer captured (*SD* = 3.4; range: 2.7–10.9) and veterinary effort averaged 2.5 hours/deer sterilized (*SD* = 2.0; range: 1.3–6.1) (Table 2). Work effort varied among sites and years. Local volunteers participated at AA, CLIF, FC, and NIH, and as appropriate, volunteers handled tasks such as pre-baiting, site preparations, transportation, and surgery when qualified. We could not accurately account for volunteer hours and our time spent training and providing oversight to volunteers. We used effort per deer and not total person-hours, which included pre-capture preparations and travel. Annual increases in capture effort per deer occurred as fewer untreated deer were available in each study area. The total cost for Year 1, including all projects, was $434,440, with 423 deer captured and 368 female deer sterilized (Table 3). If FC Year 1 is removed from the analysis because VA DGIF did not permit bait use, then the average cost per deer sterilized was $1,185.

**Table 2.** Field staff and veterinary effort for 5 surgical sterilization study sites. We did not track staff time in AA. Hours accrued for veterinarians are summarized as veterinary person-hours/deer sterilized (VT H/S), and hours accrued for field staff are summarized as field staff person-hours/deer captured (FS H/C). In CLIF and AA, volunteers participated in pre-baiting, site preparations, surgery, and transport of animals. Volunteers participated in pre-baiting and site preparations at FC. Volunteers participated in surgeries at NIH. Volunteer hours are not included, but overall represent minimal impact given the training and support needed to train volunteers. CLIF = Clifton area, Cincinnati, Ohio, USA; FC = City of Fairfax, Virginia, USA; NIH = National Institutes of Health, Bethesda, Maryland, USA; VCH = Village of Cayuga Heights, New York, USA; VGCC = Villages Golf and Country Club, San Jose, California, USA

|  | Year 1 |  | Year 2 |  | Year 3 |  | Year 4 |  | Year 5 |  |
| --- | --- | --- | --- | --- | --- | --- | --- | --- | --- | --- |
| Study |  |  |  |  |  |  |  |  |  |  |
| sites | VT H/S | FS H/C | VT H/S | FS H/C | VT H/S | FS H/C | VT H/S | FS H/C | VT H/S | FS H/C |
| CLIF | 1.6 | 3.0 | 3.3 | 6.6 | 6.8 | 9.8 | 9.1 | 13.3 | NA | 15.0 |
| FC | 6.1 | 10.9 | 6.8 | 12.3 | 5.2 | 7.7 | 5.0 | 9.8 | 5.0 | 10.1 |
| NIH | 2.0 | 4.3 | 2.2 | 4.4 | 1.5 | 2.8 | 2.0 | 3.0 | 1.3 | 2.7 |
| VCH | 1.3 | 3.7 | 2.2 | 9.1 |  |  |  |  |  |  |
| VGCC | 1.6 | 2.7 | 6.0 | 9.0 |  |  |  |  |  |  |
| Mean | 2.5 | 4.9 | 4.1 | 8.3 | 4.5 | 6.8 | 5.4 | 8.7 | 3.2 | 9.3 |

**Table 3.**
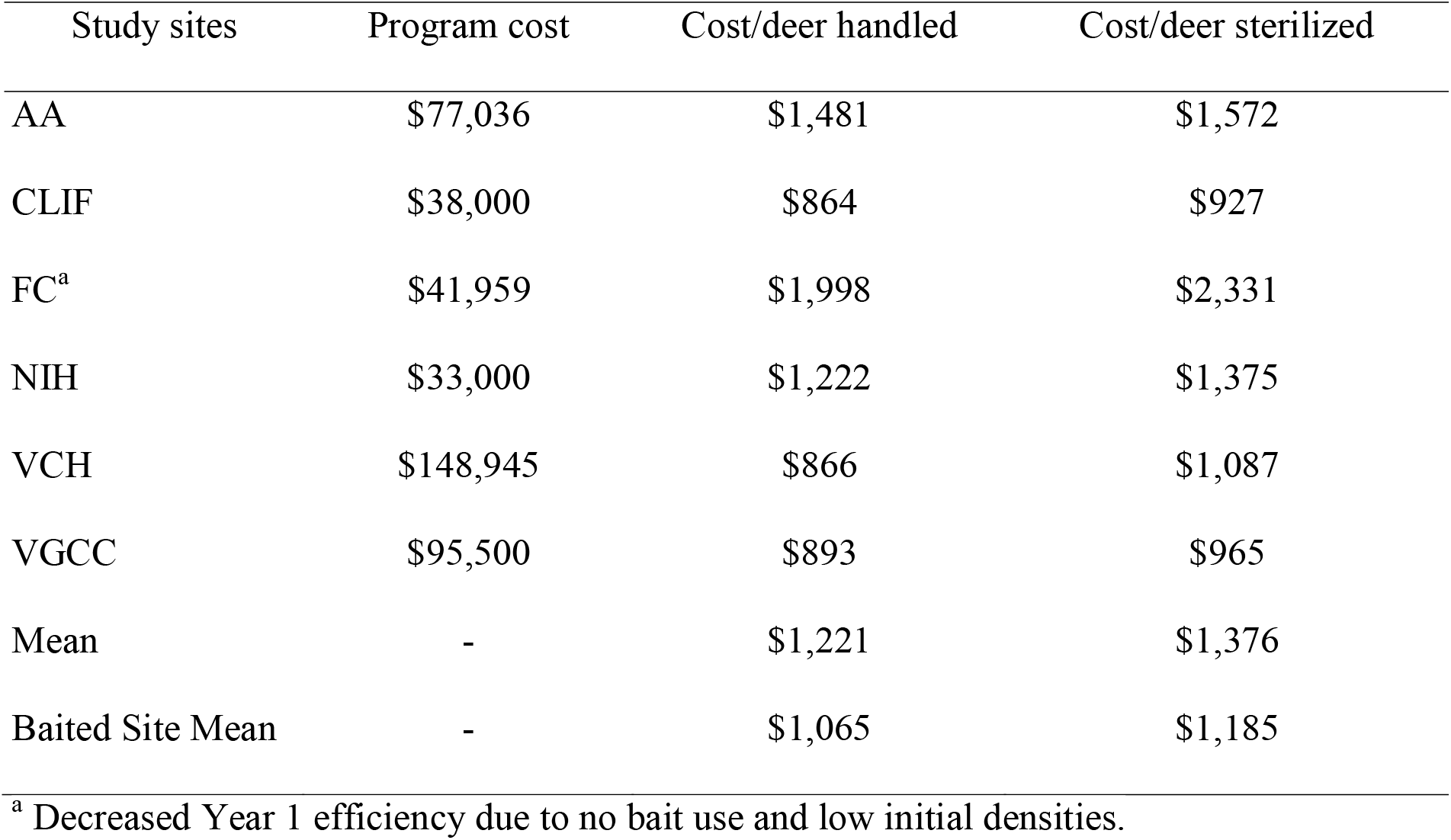
Cost per white-tailed deer handled and cost per deer sterilized in Year 1 at 6 surgical sterilization study sites. AA = Ann Arbor, Michigan, USA; CLIF = Clifton area, Cincinnati, Ohio, USA; FC = City of Fairfax, Virginia, USA; NIH = National Institutes of Health, Bethesda, Maryland, USA; VCH = Village of Cayuga Heights, New York, USA; VGCC = Villages Golf and Country Club, San Jose, California, USA

| Study sites | Program cost | Cost/deer handled | Cost/deer sterilized |
| --- | --- | --- | --- |
| AA | \$77,036 | \$1,481 | \$1,572 |
| CLIF | \$38,000 | \$864 | \$927 |
| FC<sup>a</sup> | \$41,959 | \$1,998 | \$2,331 |
| NIH | \$33,000 | \$1,222 | \$1,375 |
| VCH | \$148,945 | \$866 | \$1,087 |
| VGCC | \$95,500 | \$893 | \$965 |
| Mean | - | \$1,221 | \$1,376 |
| Baited Site Mean | - | \$1,065 | \$1,185 |

<sup>a</sup> Decreased Year 1 efficiency due to no bait use and low initial densities.

### Population Estimates and Immigration

For all 6 sites combined, we noted a mean reduction in deer abundance of about 25% (range: 16.2%–36.2%) from Year 1 to Year 2 (Table 4, Table 5). Four years after the first treatment at sites managed with only surgical sterilization (CLIF, FC, NIH, VGCC), we noted a mean total population reduction of 45% (range: 28%–56%), and a mean reduction of 90% in fawn to adult female ratio. Overall, we demonstrated annual population declines of 3–36%, with an average annual reduction of 16%. This does not include the last two years at NIH where very small changes in abundance with few deer remaining resulted in relatively large percentage swings (e.g., −8.7%, 44%). We detected 85 antlerless immigrants across all study sites between 2012– 2020, with 52 immigrant individuals detected at VCH. From 2015–2018 in VCH, 130 deer were culled using archery and remote immobilization equipment resulting in a decline from 137 to 9 deer (Table 6).

**Table 4.**
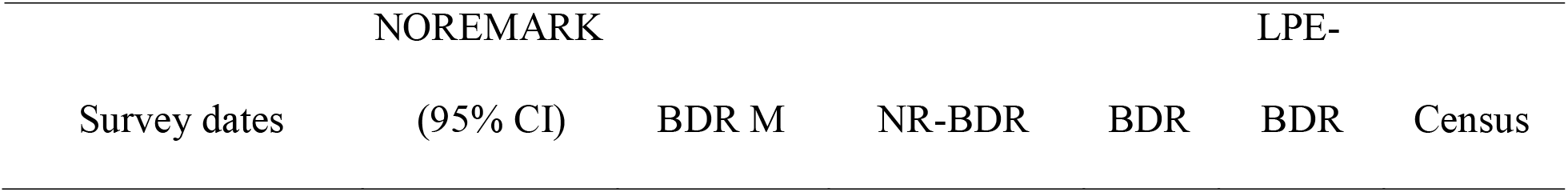
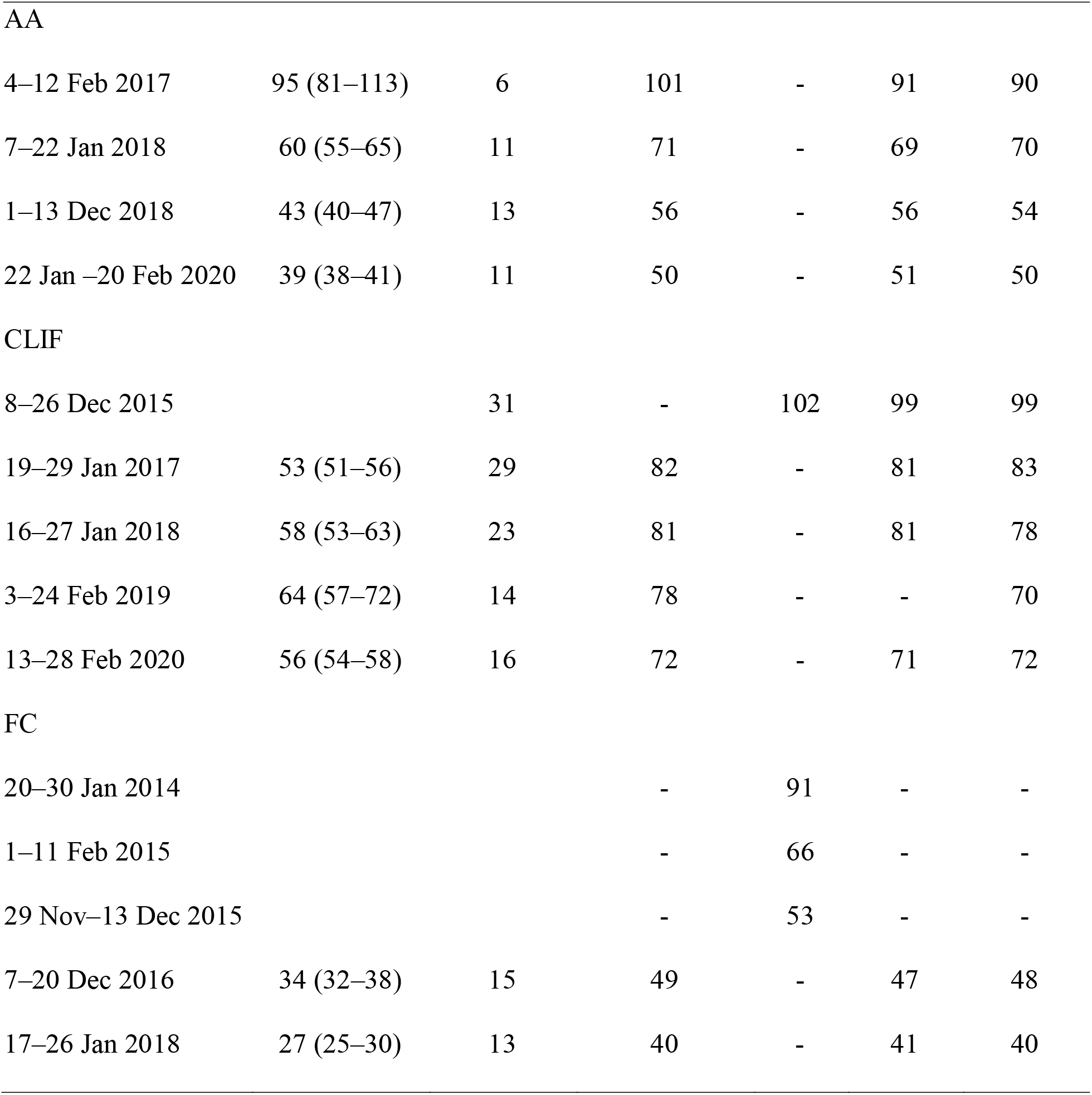
White-tailed deer population estimates at 3 study sites derived by NOREMARK, BDR, LPE and Census. The final estimate was an average of the methods. NR-BDR = NOREMARK estimate for antlerless deer and antlered males found using BDR method; LPE-BDR = LPE method for antlerless and antlered males found using BDR method; AA = Ann Arbor, Michigan, USA; CLIF = Clifton area, Cincinnati, Ohio, USA; FC = City of Fairfax, Virginia, USA

**Table 5.**
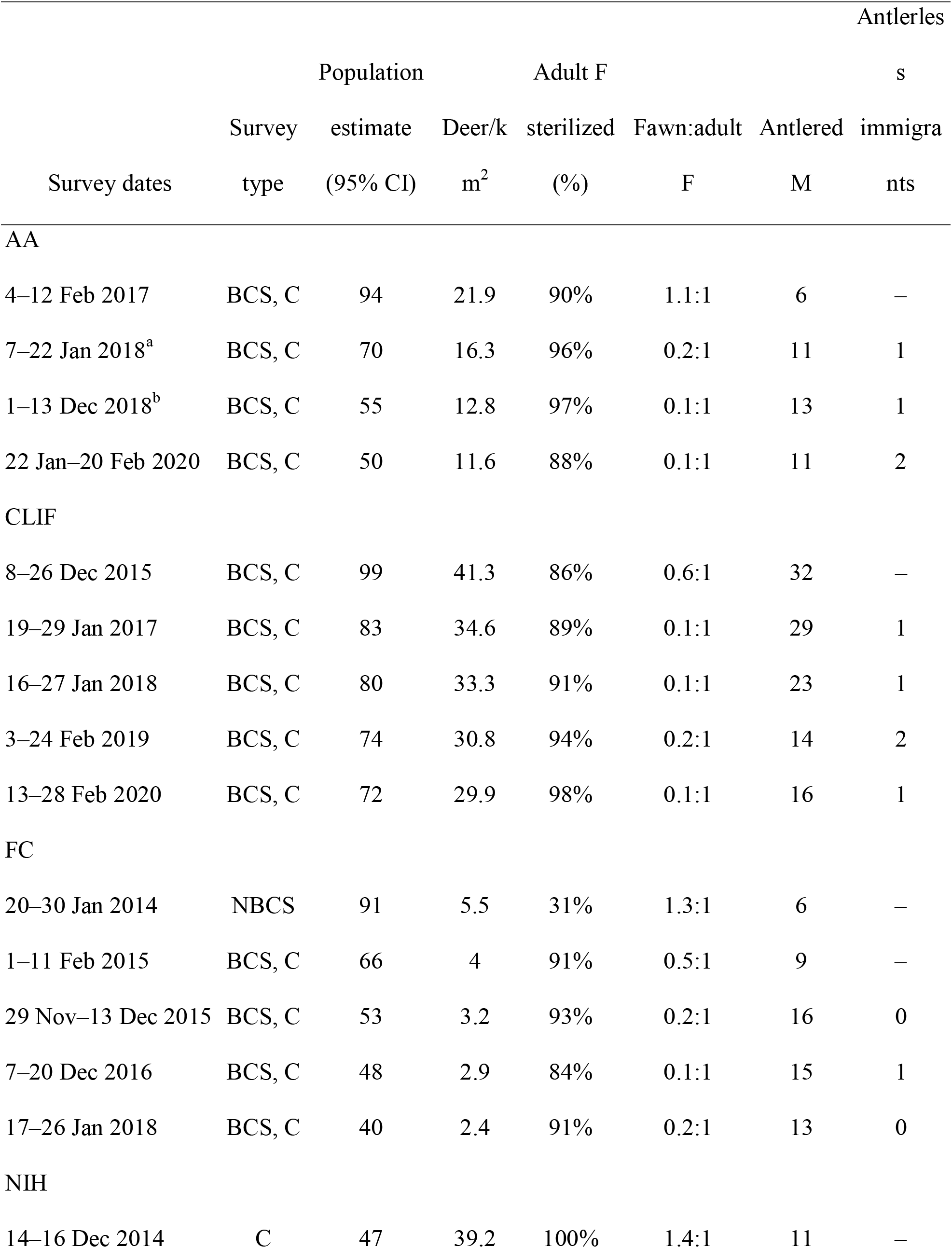
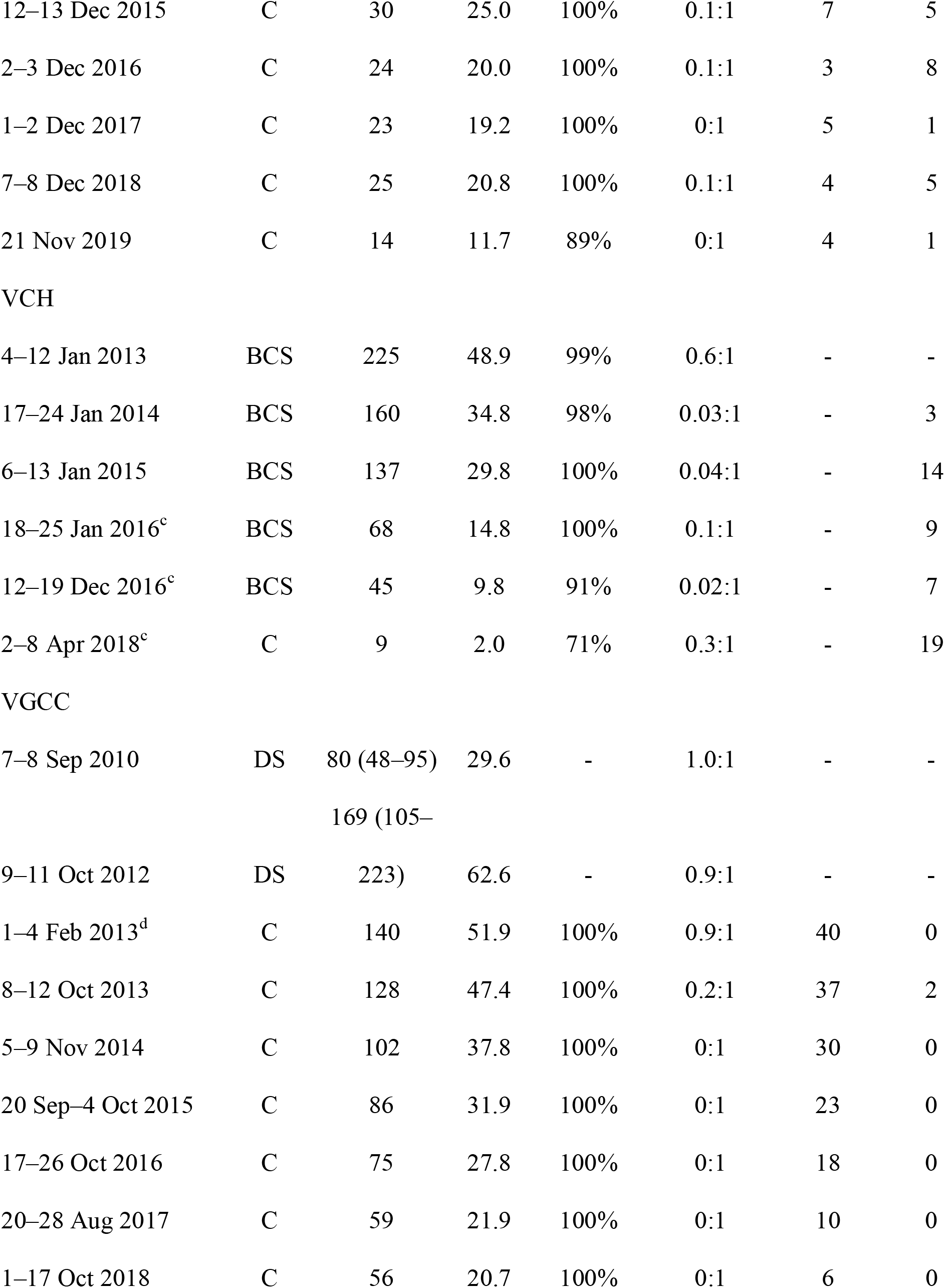
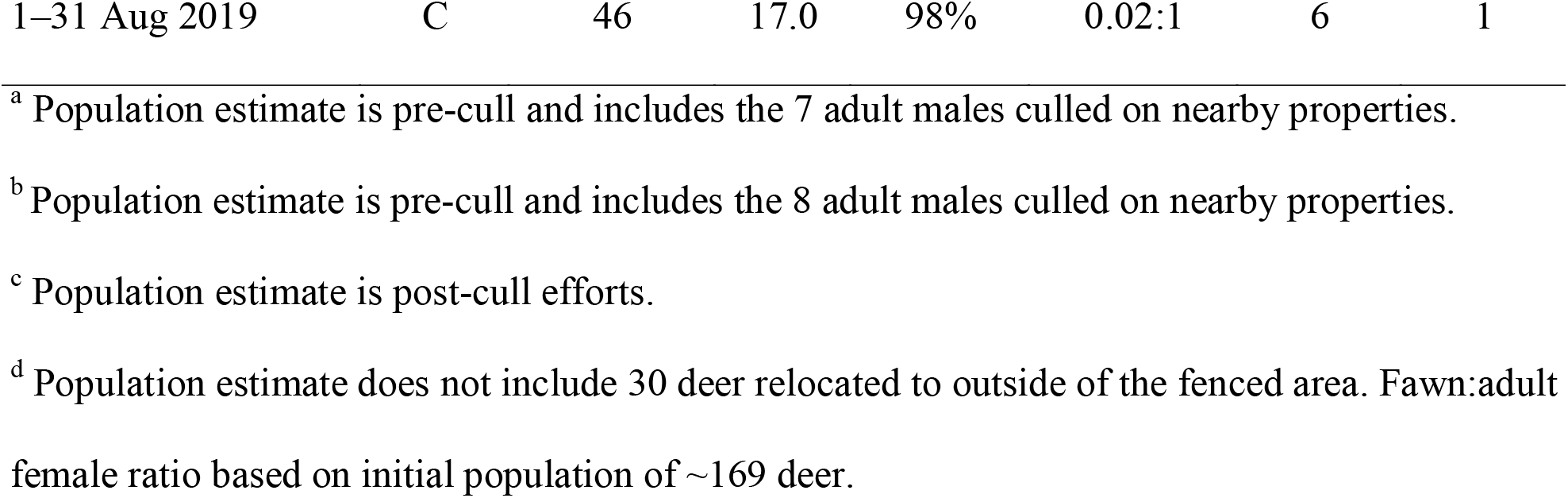
White-tailed deer population demographics derived from population estimates and field operations for 6 surgical sterilization study sites from 2012–2020. BCS = Baited camera survey; C = Census; DS = Distance sampling; NBCS = Non-baited camera survey; AA = Ann Arbor, Michigan, USA; CLIF = Clifton area, Cincinnati, Ohio, USA; FC = City of Fairfax, Virginia, USA; NIH = National Institutes of Health, Bethesda, Maryland, USA; VCH = Village of Cayuga Heights, New York, USA; VGCC = Villages Golf and Country Club, San Jose, California, USA

**Table 6.**
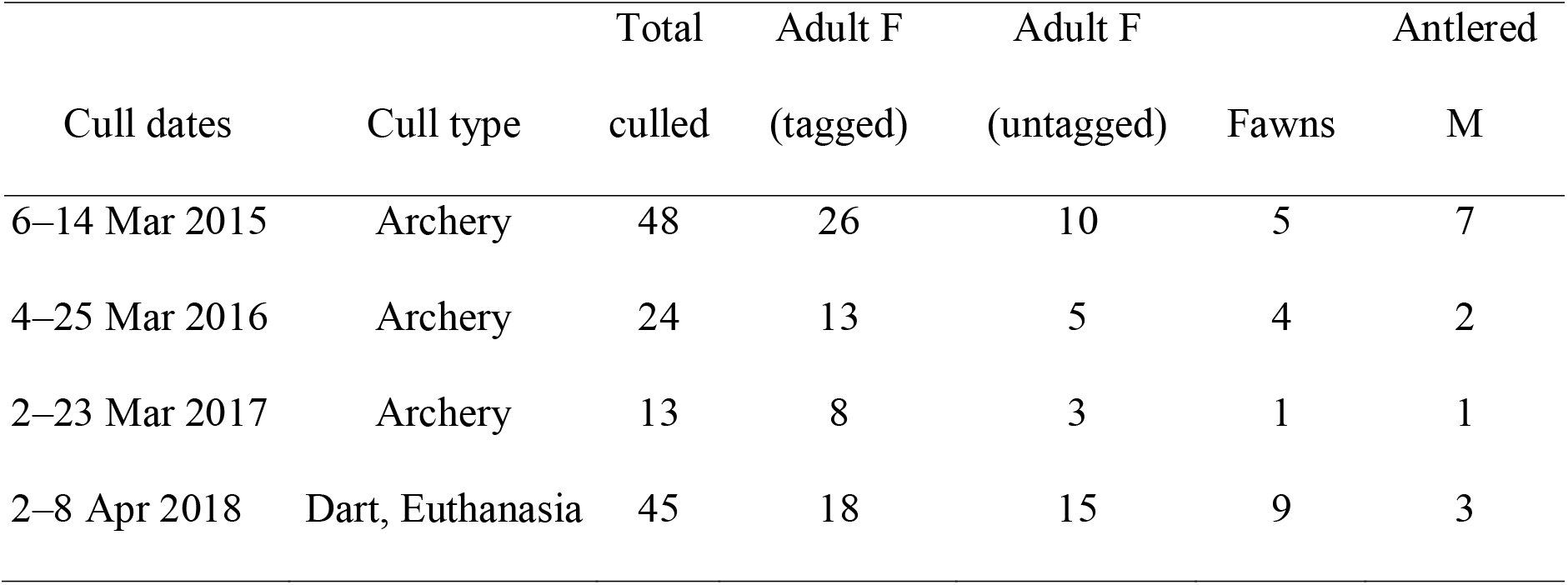
Annual white-tailed deer cull data segregated by tag status, sex, and age at Village ofCayuga Heights, New York, USA, from 2015 to 2018.

## DISCUSSION

Improvements in efficacy and longevity of non-surgical wildlife infertility methods have stagnated, and under some circumstances, surgical sterilization has been shown to be more cost-effective with longer-term efficacy (Evans et al. 2016). Cost for vaccine treatments exceeded $700, demonstrating high costs, >20 years ago (Curtis et al. 1998) with minimal improvements in vaccine administrative methods since that time. We documented that the average price per deer treated using a surgical approach and bait was $1,185, similar to the amount reported by others (Boulanger and Curtis 2016, Mathews et al. 2005). Present vaccine treatment options have changed minimally since early summaries of fertility control technology (Bomford 1992, Warren 1995, Miller et al. 1998). Reduced efficacy to ~50% after one year with single-dose vaccines is still a significant limitation for both gonadotropin-releasing hormone-based and porcine zona pellucida-based preparations (Gionfriddo 2011, Rutberg et al. 2013a). Therefore, we concluded that the permanence and 99% effectiveness of surgical methods (MacLean et al. 2006) were necessary to assess localized impacts of deer management via fertility control.

We have demonstrated that permanently sterilizing ≥85% of female deer in 6 study sites resulted in average annual population declines of 16%, with the largest local population decline occurring between Year 1 and Year 2. This 25% average decline between Year 1 and Year 2 was the result of mortality, a substantial reduction in fawn recruitment, and dispersal of male fawns as they aged to yearlings. There were few yearling males present during Year 2 capture efforts, relative to the abundance of male fawns during Year 1 capture efforts, reflecting that a high percentage dispersed the spring and fall after the first field session. Given nearly 20% of non-hunted population are composed of male fawns (DeNicola et al. 2008), this behavior contributed substantially to the initial population reduction. This level of reduction is contrary to most previous study projections when using fertility control treatments for managing deer in geographically open environments (Seagle and Close 1996, Merrill et al. 2006, Boulanger et al. 2012). Other fertility control programs in geographically closed island or fenced environments have documented similar rates of annual population decline (Naugle et al. 2002, Rutberg and Naugle 2008, Rutberg et al. 2013b). Of our study sites, only NIH and VGCC had security fencing, and neither fence prevented deer from entering or exiting. Our tagged and VHF collared deer at VGCC were often observed outside of the property. We reduced the local deer population by ~73% over 7 years at VGCC by sterilization alone. The program has successfully met community’s deer population goals, and some residents are now concerned that the population will be reduced too low (A. DeNicola, personal communication). This is an improbable outcome based on previous expectations of fertility control methods for managing deer populations. Nearly all females need to be treated in populations with annual survival rates >90% to achieve a population decline (Grund 2011). With nearly 90% treatment rates, we obtained about an 8% population reduction from Year 3 to Year 4 in CLIF, and about 28% over 4 years, where we documented 2 VHF collared female mortalities out of 17 collared over the first 3 years.

We selected research locations based on the fundamentals that emphasized the need for extensive access and approachable deer (Garrott 1995, Rudolph et al. 2000). Mobile darting from a vehicle on public roads and use of bait was critical in these case studies. By using an approach method that deer were exceptionally accustomed to, we were able to capture nearly all the females before they fully recognized the threat and began to avoid our capture efforts (Kilpatrick et al. 1997). If we had used vaccines that required repeated treatments and only afforded 90-95% efficacy, our ability to capture and effectively prevent conception in enough females to achieve the significant population reductions we documented would have been compromised.

We captured 74% of the total females sterilized in the first year at each study site. As a result, fawn to adult female ratios diminished rapidly after the first year of treatments, significantly changing the local herd demographics. As a result of reduced fawn recruitment, there was an initial increase in the percentage of adult males after Year 1. This change was not the result of immigration, but a proportional increase as the fawn age class diminished, increasing the percentages of both adult sex classes. We did not observe an increase in the number of adult males in any of our study sites. The abundance of adult males remained the same or decreased over time in all locations. This outcome contrasts with the Cornell University campus tubal ligation research findings. They observed a nine-fold increase in male abundance due to immigration, and therefore, a minimal population reduction (Boulanger and Curtis 2016). We demonstrated a 39% population reduction over 2 years in VCH using surgical sterilization. At each study site, we had treated at least 89% of the adult females by the second year. The reduced number of untreated deer in later years also facilitates the rapid transition to local volunteers conducting field operations, if this is a program objective.

To maintain capture efficiency when we attained high percentage capture rates, we used VHF collars intentionally placed on matriarchal females to track yet untreated females (Rudolph et al. 2000). We experienced minimal diminishing returns even after >80% of the females were treated. The collars allowed us to readily locate untagged females and maintain engagement rates. These engagement rates contrast with previous modeling projections for suburban fertility control treatments that depicted increased effort per deer treated as higher percentages are handled (Rudolph et al. 2000). Radio-collar tracking, used in conjunction with mobile darting from public roads, and the use of bait were critical to the success of all research projects. The effectiveness of mobile darting from the road was further demonstrated with all but about 9 deer lethally removed from VCH in April 2018.

Behavioral changes associated with surgical sterilization treatment via tubal ligation in Highland Park, Illinois, USA were noted with increased dispersal of treated females and increased mortality (Gilman et al. 2010). We had many female deer become unaccounted for, both collared and uncollared, but we could not always determine if they dispersed or died locally because they were never detected or recovered. We only tracked VHF collared females each year during subsequent capture sessions. We could not validate all mortalities of collared females, and therefore, could not differentiate our inability to locate collared female deer due to mortality or dispersal. Consequently, it was not possible to accurately quantify dispersal rates.

Overall, we experienced 15 mortalities while handling 570 deer, resulting in a 2.6% mortality rate for capture and post-surgery combined. NIH’s relatively high handling mortality rate resulted from training new veterinarians during the first 3 years and led to extended surgical times of 75–100 minutes. These surgical efforts were 3–4 times longer than the typical time with experienced veterinarians. There was 1 post-surgical death at NIH in Year 1, 1 in Year 2, 1 in Year 3, and none in Years 4 and 5. If we removed the NIH training-related deaths, our mortality rate was 2.0% (11/525). Acceptable mortality rates are suggested to be <2% for chemical immobilization of large mammals, so when our capture efforts are combined with a surgical procedure, we are confident in our results’ acceptability (Arnemo et al. 2006).

Although fertility control programs can be costly, there are actual costs from doing nothing as deer-related impacts increase. In Fairfield County, Connecticut, it was estimated that each household incurred an average of ~$900/year associated with DVCs, landscape damage, environmental impacts, tick treatments, and Lyme disease-associated medical costs (Arno and Viola 2010). Also, the cost of management options is relative, given that lethal methods may be severely constrained in their effectiveness in the same location because of development density (AA, VCH, VGCC), social or political resistance (FC, NIH), logistical and technical limits of archery hunting (CLIF), or legal constraints (VCH). Therefore, in situations where lethal methods are limited or not an option, fertility control methods can reduce deer densities as low or lesser than archery hunting (Williams et al. 2013) if deer are approachable by vehicles. We demonstrated deer reductions at or below the archery threshold of ~17 deer/km^2^ in AA (11.6 deer/km^2^), FC (2.4 deer/km^2^), NIH (11.7 deer/km^2^), and VGCC (17.0 deer/km^2^) (Williams et al. 2013). Longer-term data will allow us to assess whether culling or hunting in adjoining open spaces reduces immigration rates at lower densities. In AA, we documented similar population decline rates in the first few years to the other study sites that lack nearby culling programs. In some unique locations, when politics permit, the most cost-effective management approach might be a hybrid program using lethal methods in conjunction with fertility control treatments (Garrott 1995).

## MANAGEMENT IMPLICATIONS

There are an increasing number of developed areas where lethal deer management options cannot meet the stated objectives. Although not a deer management panacea, high-percentage treatment via ovariectomy can significantly reduce local deer populations, especially in areas with necessary vehicle access and approachable deer, given most legal and social obstacles are not present. As with any management technique, the project scale and relative deer abundance will affect the cost and feasibility of meeting stated objectives. Social attitudes in rural and developed landscapes align with lethal and nonlethal methods’ relative applicability, respectively. It may be necessary to incorporate lethal methods combined with this nonlethal approach, if possible, when immigration precludes meeting management objectives, or a shorter management timeline is desired. Finally, it is important to note that fertility control treatments may be a necessary co-management technique with hunting or culling programs when local populations cannot be reduced by >30-40% annually because of legal and development constraints.

## ACKNOWLEDGMENTS

We are grateful for the cooperation and permitting by the involved state wildlife agencies. We thank local law enforcement agencies for assistance and oversight. Our research would not have been possible without collaboration and support from partners such as P. Curtis from Cornell University, who was instrumental in the organizational and population estimation phases in VCH, the staff at NIH for field and surgical support, and the Cincinnati Parks Department for help in CLIF. We thank numerous veterinarians, including Dr. K. Dyer, Dr. R. Junge, Dr. J. Newman, Dr. K. Ortved, and, most of all, Dr. S. Timm, who helped refine the surgical procedure to make it both field efficient and humane. We thank the local volunteers who provided valuable field assistance. We are grateful to the many homeowners who allowed us to capture deer on their properties. Projects would not have been funded or authorized without the thoughtfulness of City Council members, local board members, and donors. Finally, we are thankful to Dr. S. Williams for his review of an earlier version of this manuscript.

Associate Editor:

## Figure Captions

**Figure 1.** Delineation of focal white-tailed deer surgical sterilization study area in Ann Arbor, Michigan, USA.

**Figure 2.** Delineation of white-tailed deer surgical sterilization study area in the Clifton area of Cincinnati, Ohio, USA.

**Figure 3.** Delineation of white-tailed deer surgical sterilization study area in City of Fairfax, Virginia, USA.

**Figure 4.** Delineation of white-tailed deer surgical sterilization study area in National Institutes of Health, Bethesda, Maryland, USA.

**Figure 5.** Delineation of white-tailed deer surgical sterilization study area in the Village of Cayuga Heights, New York, USA.

**Figure 6.** Delineation of black-tailed deer surgical sterilization study area in the Villages Golf and Country Club, San Jose, California, USA.

